# Pharmacokinetics of Orally Administered GS-441524 in Dogs

**DOI:** 10.1101/2021.02.04.429674

**Authors:** Victoria C. Yan, Cong-Dat Pham, Matthew J. Yan, Alexander J. Yan, Sunada Khadka, Kenisha Arthur, Jeffrey J. Ackroyd, Dimitra K. Georgiou, Laura E. Roon, Lane R. Bushman, Peter L. Anderson, Chun Li, Florian L. Muller

## Abstract

Despite being FDA-approved for COVID-19, the clinical efficacy of remdesivir (Veklury^®^) remains contentious. We previously pointed out pharmacokinetic, pharmacodynamic and toxicology reasons for why its parent nucleoside GS-441524, is better suited for COVID-19 treatment. Here, we assess the oral bioavailability of GS-441524 in beagle dogs and show that plasma concentrations ∼24-fold higher than the EC_50_ against SARS-CoV-2 are easily and safely sustained. These data support translation of GS-441524 as an oral agent for COVID-19.

---

Remdesivir (RDV) is currently the only FDA-approved anti-viral drug for the treatment of COVID-19 despite exhibiting just modest efficacy in one double-blind, placebo-controlled randomized clinical trial (RCT) (1); other RCTs have thus far found no statistically significant improvement in mortality (2) or time to clinical improvement (3). As a phosphoramidate prodrug of the McGuigan class (4), RDV is structurally susceptible to conversion to its active nucleoside triphosphate (NTP; GS-443902) form by enzymes that are abundant in the liver (CES1/CTSA/HINT1) but minimally expressed in alveolar type 2 cells (AT2) (5), the cell type most susceptible to SARS-CoV-2 infection (6). Preferential liver metabolism of RDV results in on-target dose limiting toxicity (DLT) that precludes dose escalation despite its modest clinical performance (7, 8). Such shortcomings are exacerbated by the hydrophobic nature of RDV, which requires complex excipients that could implicate kidney function (9, 10). Another major drawback with RDV is its requisite intravenous (IV) administration (11), making outpatient therapy, early intervention, and prophylaxis impractical.

Cognizant of these limitations, we have asserted that the parent nucleoside of RDV, GS-441524, would be better suited for the treatment for COVID-19 (5). GS-441524 is the persistent, predominant metabolite in plasma following IV infusion of RDV in preclinical species (12–14) and in humans with a half-life (T_1/2_) of >24 h (8, 15). The extent to which GS-441524 contributes to the overall anti-SARS-CoV-2 activity of RDV when administered to patients remains unclear. In contrast to RDV, which is preferentially bioconverted to GS-443902 by liver-abundant enzymes (16, 17), GS-441524 is bioconverted to GS-443902 by nucleoside kinases (likely adenosine kinase, ADK) that are broadly expressed across all tissues. Due to its demonstrably better safety profile (7, 18), a significant advantage that GS-441524 possesses over RDV is the possibility for dose escalation without liver-related DLTs, as this would increase the concentration of bioactive GS-443902 in AT2 cells. Cell-based studies have shown that GS-441524 is a potent inhibitor of SARS-CoV-2-infected cells, with EC_50_ values on the same order of magnitude as that of RDV (EC_50_= 0.47-1.09 µM) (19). Efficacy studies conducted in cats with natural presentations feline infectious peritonitis (FIP) as a result of infection by the closely related feline coronavirus (FCoV) have demonstrated up to 96% cure rate with subcutaneously (SC) administered GS-441524 (20–22). Efficacy studies conducted in mice infected with either SARS-CoV-2 or murine hepatitis virus (MHV, a closely related coronavirus) have shown that GS-441524 is capable of reducing viral loads in pathologically relevant organs without obvious adverse events (23). Given the scarcity of simple outpatient treatment options for COVID-19 (24), these especially encouraging data warrant translation of GS-441524 to the clinic. Here, we provide pharmacokinetic (PK) evidence in dogs supporting the rationale for GS-441524 to be investigated as an oral agent for COVID-19.

To validate our assertion that McGuigan prodrugs such as RDV are heavily subject to first-pass metabolism, we first conducted a single-dose, equimolar comparison between orally (PO) administered GS-441524 (6.5 mg/kg) and RDV (13 mg/kg) in dogs. Male beagles (N=1 per group) were administered excipient-less capsule formulations of either GS-441524 or RDV and plasma concentrations of GS-441524 were evaluated at predetermined timepoints (**Figure 1a**). No adverse events were observed in either dosing group. As expected, plasma concentrations of GS-441524 following administration of RDV were poor, with C_max_ values roughly 25-fold lower than that observed when GS-441524 was administered directly (172 vs. 4580 ng/mL, respectively; **Figure 1a, c, Supplementary Data File 1**). Interestingly, there was an observable difference in T_max_ values, with plasma concentrations of GS-441524 peaking at approximately 3 h following RDV administration versus 1 h following direct dosing of GS-441524. This shift in T_max_ values with RDV PO administration suggests a mechanism of systemic release similar to that observed for the McGuigan prodrug sofosbuvir, wherein rapid hepatic extraction of intact prodrug forms a reservoir of active NTP and hydrolyzed nucleoside—the latter of which is then slowly released into systemic circulation (25). Given that plasma concentrations of GS-441524 following RDV administration are below the range of reported anti-SARS-CoV-2 EC_50_ values and that long-term PO dosing of RDV at 13 mg/kg is almost certainly therapeutically prohibitive in humans due to hepatotoxicity concerns (8), these data allude to the infeasibility of administering RDV PO for COVID-19. At the same time, we find that direct PO administration of GS-441524 results in plasma concentration exceeding the range of reported anti-SARS-CoV-2 EC_50_ values for at least 8 h (**Figure 1a, Supplementary Data File 1**). Peak concentrations of GS-441524 reached 15.45 µM and were obtained at approximately 1 h (**Figure 1a, c**), indicating high oral absorption of GS-441524 in dog even in the absence of excipients.

**Figure 1.**
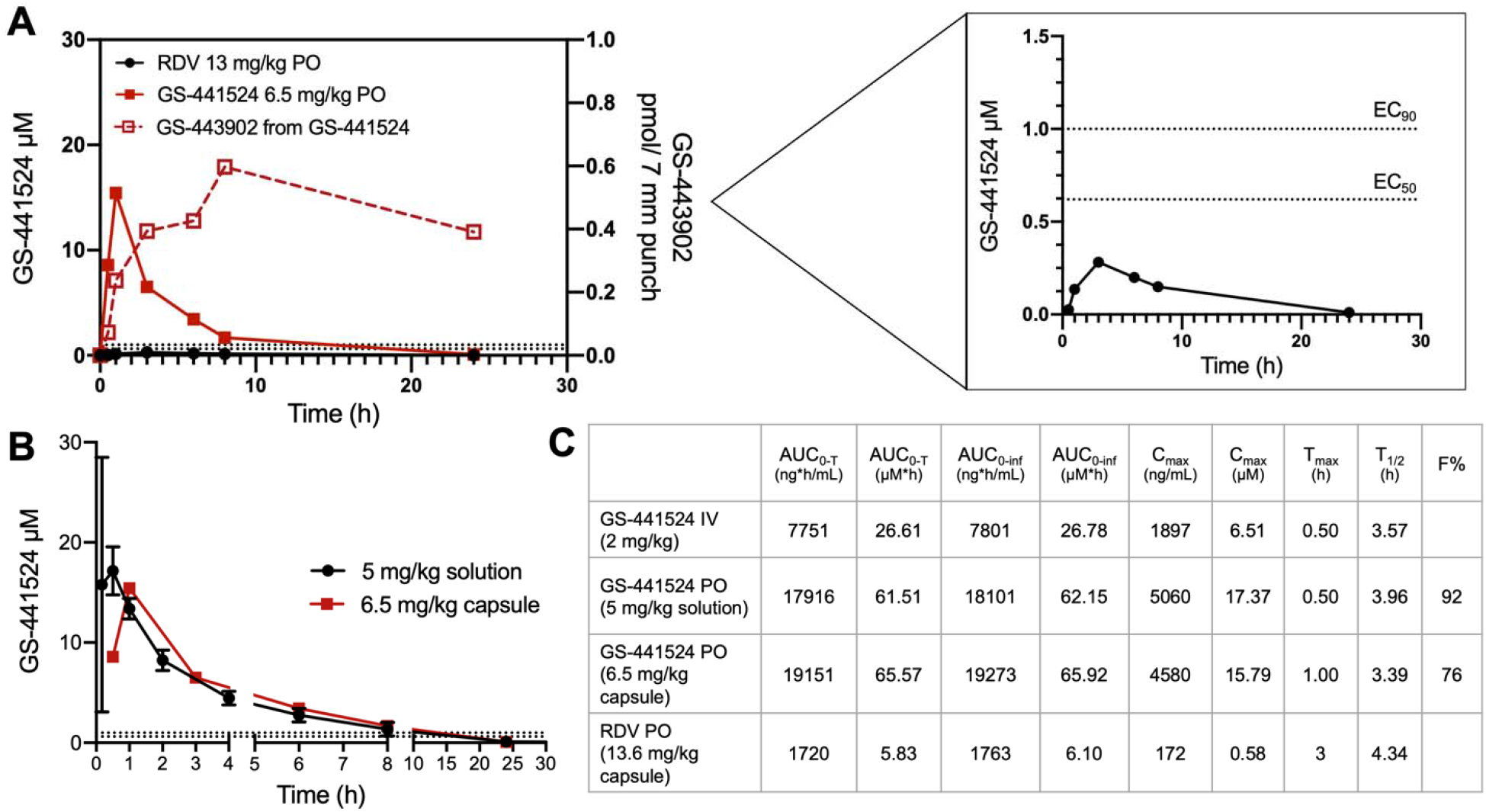
Plasma concentrations of GS-441524 following a single oral dose of remdesivir or GS-441524 in dogs. **(A)** Head-to-head PK comparison following a single equimolar dose of remdesivir (black, 13.6 mg/kg) and GS-441524 (orange, 6.5 mg/kg) in male beagle dogs (N=1 per compound). Both compounds were administered in capsule form. Plasma concentrations of GS-441524 following compound administration are plotted for the following timepoints (h): 0.5, 1, 3, 6, 8, 24. A focused view of GS-441524 concentrations following oral administration of remdesivir is shown on the left. Dashed red line corresponds to the levels of GS-443902 (NTP) formed in whole blood following oral administration of GS-441524 as quantified using a highly sensitive dried blood spot assay. Oral administration of RDV did not produce detectable levels of GS-443902 in whole blood. **(B)** Comparison of plasma concentrations of GS-441524 following oral administration as a solution (black, 5 mg/kg; N=3) or as a capsule (orange, 6.5 mg/kg; N=1). **(C)** Mean PK parameters following various routes of administration of GS-441524 and RDV. Raw values for GS-441524 dosed IV and PO dosed as a solution are adapted from NCATS OpenData Portal and have been re-calculated to match the sampling timeframe of the capsule studies (T=0.5-24 h). All PK parameters were calculated using PKSolver 2.0. In panels A and B, dotted lines correspond to EC_50_ (bottom) and EC_90_ (top) values reported for GS-441524 in SARS-CoV-2-infected Calu3 cells (19).

Prior studies assessing the oral bioavailability (F%) of GS-441524 in dogs following both IV (2 mg/kg) and PO (5 mg/kg) administration have found that the drug is efficiently absorbed, with an F% of 85% (NCATS OpenData Portal). Such studies examined PO absorption of GS-441524 using a solution formulation prepared at a final concentration of 2.5 mg/mL. We sought to determine whether similarly favorable F% could be achieved using an excipient-less capsule formulation, which would greatly ease outpatient administration. The wide range of F% observed in other preclinical species (**Table 1**; NCATS OpenData Portal) and the unusual solubility properties of GS-441524 are reminiscent of that observed with the FDA-approved nucleoside analogue acyclovir (**Table 1**), which was ultimately formulated as an excipient-less tablet (26). Direct comparison of PK parameters between solution and capsule formulations of GS-441524 indicates a similar pattern of high drug absorption, with T_max_ values of 0.5 and 1 h, respectively (**Figure 1b**). Between solution and capsule formulations, PK parameters were generally similar; it should be noted that the capsule dose was slightly higher than the solution dose (5 mg/kg vs. 6.5 mg/kg). Adjusting for sampling timeframes, these data indicate that the C_max_ value was higher with the solution formulation (5060 vs. 4580 ng/mL) but the AUC value were somewhat higher with the capsule formulation (17,916 vs.19,151 ng*h/mL; **Figure 1c**). Such observations appear consistent with the general observation that liquid formulations tend to be more readily absorbed than their pill counterparts (27). Nevertheless, the estimated F% using this capsule formulation remains high at about 76% (**Figure 1b**). Pharmacodynamic comparison GS-441524 and RDV corroborates the ability for GS-441524 but not RDV to be administered orally. Quantification of NTP formation in whole blood using a highly sensitive dried blood spot assay found that plasma concentrations of GS-441524 following direct administration was able to generate levels of GS-443902 in whole blood; in contrast, GS-441524 produced as a metabolite of orally administered remdesivir was not able to form levels of NTP above the lower limit of quantification (**Figure 1a, dashed line**). These data hint at the feasibility of using an excipient-less pill formulation for GS-441524 for outpatient treatment.

**Table 1.** Oral bioavailabilities of acyclovir and GS-441524 in preclinical species are similar. GS-441524 exhibits a similar pattern of F% as acyclovir across preclinical species. F% for GS-441524 were obtained from the NCATS OpenData Portal and F% for acyclovir were obtained from the FDA fact sheet on Zovirax (human), Laskin et al. Clin. Pharm. (1983, monkey) (35) and Krasny et al. *J. Pharm. Exp. Ther*. (1981, dog) (36). *Anticipated F% of GS-441524 in humans.

| Species, F% | Acyclovir | GS-441524 |
| --- | --- | --- |
| Human | 10-20 | 15-30* |
| Monkey | 3.7 | 8.3 |
| Dog | 54-90 | 85-93 |

There are some limitations associated with this study. First, the sample size in the capsule study is small, which may not capture possible variability associated with this formulation. Second, nucleoside analogues generally tend to exhibit higher F% in dogs than in other preclinical species perhaps due to the presence of a paracellular nucleoside transporter that is absent in humans and non-human primates (28). As a result, the F% of nucleoside analogues in dogs tends to overestimate that observed in humans (**Supplementary Data File 2**). While not specifically explored in this study, it should be noted that—at the other end of the F% spectrum—F% of nucleoside analogues in non-human primates tend to under-predict that observed in humans (**Supplementary Data File 3**). Nevertheless, these data suggest the feasibility of using a simple, excipient-less capsule formulation of GS-441524. As a prodrug inhibitor of the SARS-CoV-2 RNA-dependent RNA polymerase (RdRp), GS-441524 is aptly poised to demonstrate consistent efficacy among new mutations of SARS-CoV-2, as RdRp is much less susceptible to efficacy-altering mutations than is the spike protein (29, 30). Thus, clinical translation of GS-441524 is imperative to human health.

## Drug formulation

For capsule studies, GS-441524 and RDV were purchased at the highest commercially available quality from MedKoo Biosciences; purity was verified by ultra-performance liquid chromatography mass spectrometry (UPLC-MS) and nuclear magnetic resonance (NMR) spectroscopy (^1^H, ^13^C) in-house. For capsule studies, gelatin capsules (size 5, XPRS Nutra) were tightly packed with either GS-441524 (65 mg) or RDV (136.74 mg) without additional excipients. For solution studies, GS-441524 was purchased at the highest commercially available from AK Scientific and characterized by NCATS. Formulations for PO and IV studies conducted by NCATS are described on the NCATS OpenData Portal. Briefly, GS-441524 was dissolved in a solution containing 5% ethanol, 30% propylene glycol, 45% PEG-400, 20% water with 1 equivalent HCl for a final concentration of 2.5 mg/mL.

## Single dose GS-441524 and RDV in dogs via capsule formulation

All capsule form studies were performed at Charles River Laboratories (Wilmington, MA) with IACUC approval (#20236536). Fasted male adult beagles (10 kg; N=1 per compound) were administered either GS-441524 (6.5 mg/kg) or RDV (13 mg/kg). Plasma samples were taken for PK analysis at the following timepoints (h): -0.5, 0.5, 1, 3, 6, 8, 24. Animals were monitored continuously by veterinarians for any clinically relevant abnormalities during dosing and sample collection. PO solution and IV studies were performed by NCATS as described on the NCATS OpenData Portal with relevant committee and regulatory approval. Data from all studies were (re-)analyzed using PKSolver 2.0 and graphs were generated using GraphPad Prism 8.

## Plasma pharmacokinetics

For capsule studies, plasma levels of GS-441524 were analyzed at Covance, Inc (Princeton, NJ) on a fee-for-service basis using a liquid chromatography mass spectrometry (LC-MS) assay previously described for quantification of GS-441524 following IV administration of RDV in NHP (12).

## Quantification of GS-443902 in dried blood spots

Analyses were performed at CU Anschutz adapted from previously published procedures (31–34). The assay range was 0.1 – 100 pmol/sample, with a lower limit of quantitation of 0.1 pmol/sample. A 7mm disc was punched from the DBS and extracted with two mL of 50:50 methanol:water of which, 1.6 mL was assayed (this was the sample). Results were then normalized to pmol per 7mm punch.

## Author Contributions

V.C.Y. and F.L.M. conceived of the study. V.C.Y. analyzed data and wrote the manuscript. C.D.P. prepared capsule formulations of GS-441524 and RDV. L.R.B., L.E.R., and P.L.A. performed pharmacodynamic analyses on dried blood spots. M.J.Y., A.J.Y., S.K., K.A., D.K.G., and J.J.A. provided technical assistance. V.C.Y., F.L.M. and C.L. oversaw the study.

## Acknowledgements

This work was supported by the COVID-19 Early Treatment Fund (F.L.M.) and the NIH (R21AI159246-01; C.L.). S.K. is supported by the MD Anderson CPRIT Research Training Program Grant (RP170067) and the Larry Deaven PhD Fellowship in Biomedical Sciences.

